# cDNA-detector: Detection and removal of cDNA contamination in DNA sequencing libraries

**DOI:** 10.1101/2021.08.11.455962

**Authors:** Meifang Qi, Utthara Nayar, Leif S. Ludwig, Nikhil Wagle, Esther Rheinbay

## Abstract

Exogenous cDNA introduced into an experimental system, either intentionally or accidentally, can appear as added read coverage over that gene in next-generation sequencing libraries derived from this system. If not properly recognized and managed, this cross-contamination with exogenous signal can lead to incorrect interpretation of research results. Yet, this problem is not routinely addressed in current sequence processing pipelines. Here, we present cDNA-detector, a computational tool to identify and remove exogenous cDNA contamination in DNA sequencing experiments. We apply cDNA-detector to several highly-cited public databases (TCGA, ENCODE, NCBI SRA) and show that contaminant genes appear in sequencing experiments where they lead to incorrect coverage peak calls. Our findings highlight the importance of sensitive detection and removal of contaminant cDNA from NGS libraries before downstream analysis.

## Introduction

Massively parallel DNA sequencing experiments including ChIP-seq, ATAC-seq, whole exome and whole genome sequencing are widely used research methods. These methods have been instrumental for identifying binding sites of DNA-associated proteins, histone modification states, chromatin accessibility, and germline and somatic genomic variants. In common functional studies, a wild-type gene or variant of interest is introduced into the experimental system with DNA vectors, which can then appear in the sequencing library. Sequence reads stemming from the exogenous gene of interest are then mapped back to the endogenous locus during the alignment step. In addition, trace amounts of vector DNA from other experiments performed in the same laboratory or on shared equipment can contaminate the DNA library, leading to the presence of foreign genes in the derived alignment. Signal from these contaminant genes can then affect downstream results and interpretation, for example through spurious increased copy number (Kim et al., 2020; Lee et al., 2018), false germline or somatic variant calls (Kim et al., 2016; Lim et al., 2015), or incorrect coverage peak calls (Corces et al., 2018; Kim et al., 2016; Lim et al., 2015).

Detection of such exogenous gene signal in an alignment can be challenging, in particular when the vector-derived signal overlaps true signal derived from exome or chromatin profiling. Few methods have been developed to detect vector contamination in NGS libraries. Some identify sequence reads from known cloning vectors, but do not search for the cDNA insert (https://sourceforge.net/projects/seqclean/, https://www.ncbi.nlm.nih.gov/tools/vecscreen/). Vecuum (Kim et al., 2016) was developed to also identify the contaminating gene; however, this tool relies on known vector backbone sequences, and paired-end reads are required.

Because cloned cDNAs do not contain introns or their physiologic UTR regions, this property can be exploited to identify potential contaminants. Reads from these cDNAs only partially align to the genome at exon boundaries, with unmapped sequence matching either a vector (at the 5’ and 3’ ends) or a neighboring exon. In a genomic alignment, these reads appear as “clipped”, and thus can be distinguished from true signal reads that fully map to the genome across exon boundaries (**Figure 1A**).

**Figure 1:**
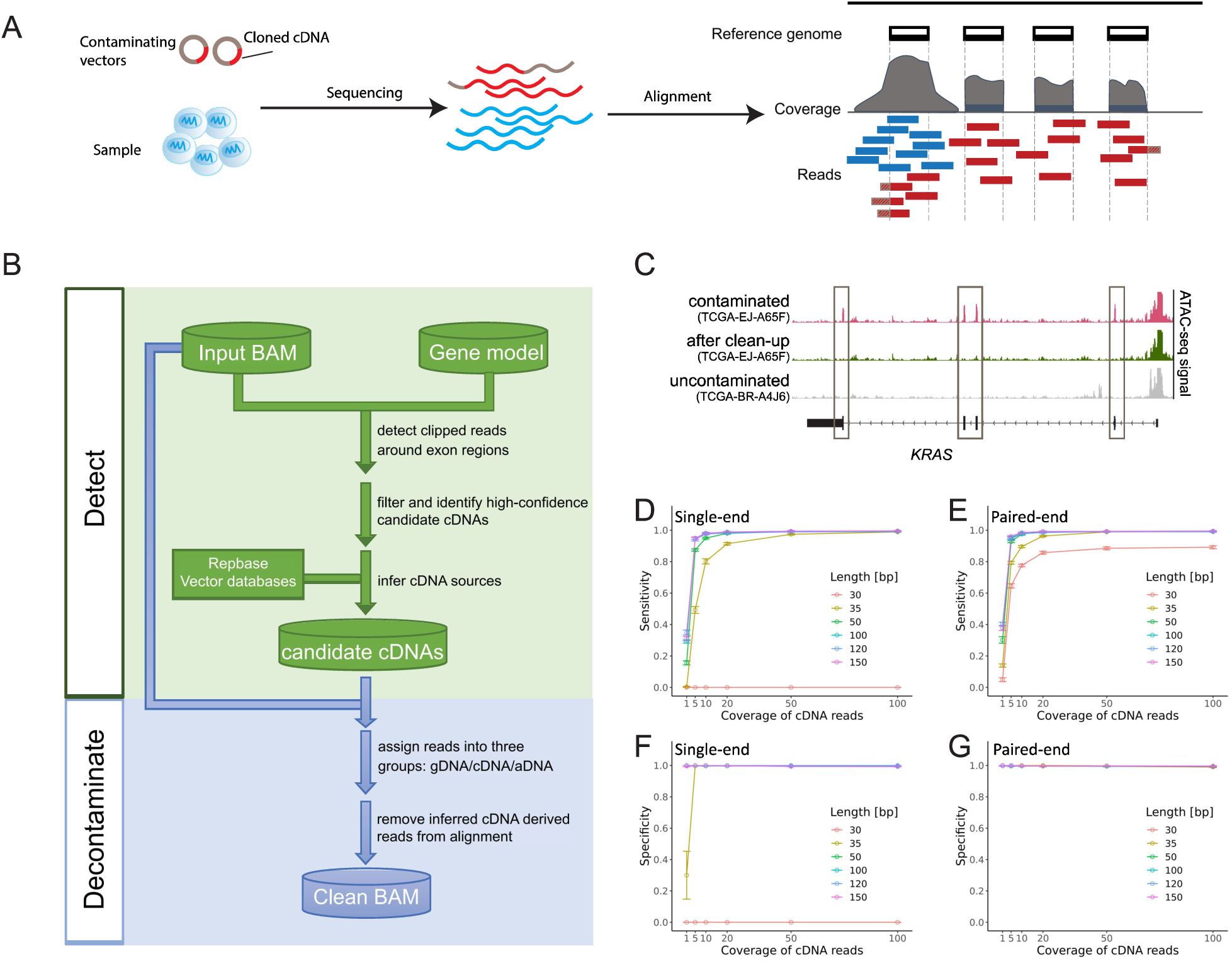
Workflow and performance of cDNA-detector. **A**, Schematic illustrating the source of intentional or contaminant cDNA reads in sequencing experiments. Vectors (grey) with cDNA (red) are amplified in the library preparation process and sequenced together with the experiment. Upon alignment of reads, cDNA-derived reads (red boxes) map to the respective gene locus in the genome, along with true signal reads (blue). Textured red read segments indicate sequence not mapping to the genome and “clipped” by the alignment algorithm. **B**, Overview of the two main components of the cDNA-detector algorithm, “detect” and “decontaminate”. **C**, ATAC-seq experiment from TCGA before (pink) and after (green) removal of vector-induced *KRAS*. For comparison, an uncontaminated sample (grey) is shown. Boxes indicate contaminant signal over exons. **D**-**E**: Sensitivity of cDNA-detector on single-end (**D**) and paired-end (**E**) sequencing experiments, depending on read coverage. **F-G**: Specificity of cDNA-detector on single-end (**F**) and paired-end (**G**) read data dependent on coverage. Error bars indicate standard errors. Random sample size n = 10 for each library type and read length.

## Results

### Detection of cDNA in NGS libraries

Prompted by the presence of intentional and accidental cDNA cross-contamination in our own data, we developed cDNA-detector, a computational tool that detects and optionally removes cDNA contamination in DNA sequencing data (**Figure 1B**). cDNA-detector takes a standard BAM-formatted alignment and a gene model file as input. For each exon from the model file, the number of clipped and properly mapped reads at the two boundaries (the start and end coordinate) are counted. Using a binomial model, we test whether the fraction of clipped reads at any given boundary coordinate exceeds the expectation based on the background of total clipped reads (Methods). The two *P*-values for each exon and all *P*-values for each transcript are combined. Significant transcripts (*P*<0.05) or transcripts with more than 30% of significant exons are considered candidate contaminant transcripts for further analysis (**Figure S1**).

Next, cDNA-detector performs an in-depth search for additional evidence at exon boundaries for all exons in candidate contaminant transcripts. First, it seeks clipped reads within a range of 5 bp of the annotated exon boundary (**Figure S1**). This step captures clipped reads when part of the clipped region matches the adjacent intronic sequence, and thus the clip is not performed at the exact boundary coordinate; and it allows for additional space for potentially added sequence at the cDNA ends that could have been introduced during cloning. Further, cDNA-detector infers a consensus sequence from the clipped overhangs. It then searches the alignment for additional reads with short overhangs (1-2 bp) annotated as mismatches rather than clips, but whose overhang sequence aligns to the inferred consensus sequence at the respective boundary (**Figure S1**). Additional evidence from reads identified in these steps is added to the clipped read count, and *P*-values are re-calculated for all candidate contaminant exons and transcripts as described above. cDNA-detector further checks whether the consensus sequence of the clipped region maps to the neighboring exon.

### cDNA source inference

Eukaryotic genomes contain endogenous retrogenes, processed (spliced) copies of genes that were reverse transcribed and re-inserted into the genomic sequence, often flanked by LINE-1 elements in the human genome (Esnault et al., 2000; Kaessmann et al., 2009; Wei et al., 2001). Similar to contaminant cDNAs, these retrogenes are also intron-free and generate clipped reads at the source gene locus, and thus cDNA-detector will detect these instances. While retrogenes are rarely picked up in ChIP-seq and ATAC-seq data, they can dominate genomic sequencing strategies (whole exome and whole genome). The tool therefore attempts to distinguish vector-induced contaminants from retrogene copies (Methods; **Figure S1**). To infer the source of a given cDNA candidate, a database of known cloning vectors (https://www.ncbi.nlm.nih.gov/tools/vecscreen/univec/) as well as consensus sequences of transposable elements (Bao et al., 2015) are queried for the consensus sequences derived from clips at the 5’ and 3’ ends of candidate transcripts. In the common case where clipped sequences are too short to yield a database query result, we consider the distance of the clipped position to the annotated boundary: if the distance is ≤ 5 bp, a vector is likely the source; if the distance is >10 bp, the cDNA candidate is deemed more likely to originate from a retrogene insertion. Cases where no clear evidence can be found (e.g. custom plasmids, short sequences) are annotated as “unknown”, and the user is encouraged to manually review potential sources. When using cDNA-detector on genomic sequence data, we recommend suppressing the “retrocopy” output, such that only potential vector cDNA candidates are reported.

### Alignment decontamination

If potential contaminants are identified, cDNA-detector provides the option to remove contaminant reads from the alignment **(Figures 1B and S2A-B**) to reduce the risk of spurious coverage peak and variant calls in downstream analysis. In addition to obvious contaminant clipped reads as identified above, cDNA-detector further classifies all reads in candidate cDNAs as either genomic or ambiguous (**Figure S2B**; Methods). Genomic reads span the exon boundary and properly map to the adjacent intron sequence; ambiguous reads do not overlap the boundary but instead map entirely inside the exon, and could thus originate from either true signal or the contaminant. In the decontamination step, cDNA-detector removes all *bona fide* contaminant reads. For ambiguous reads, the tool calculates the fraction of contaminant-to-genomic reads at the exon boundaries, and randomly discards the same fraction of ambiguous reads. This approach ensures that any true signal obscured by cDNA contamination remains in the decontaminated sample. With this strategy, contaminants can be effectively removed from alignments, revealing true signal previously obscured by contamination (**Figures 1C and S2C**).

### cDNA-detector sensitivity and specificity

We evaluated cDNA-detector’s sensitivity and specificity by generating sets of simulated cDNAs with different read lengths, distinct sequencing strategies (single vs. paired-end) and variable coverage, and introducing these into a contamination-free ATAC-seq experiment (2x 38 bp reads, 43,344,211 reads total; Methods). We found high sensitivity (>80%) for detecting contaminants when using read lengths above 50 bp and coverage greater than 5x (**Figures 1D-1E**). However, cDNA-detector failed on very short, single-end reads because the aligner (Li and Durbin, 2009) with default parameters does not produce clipped reads when the read length is ≤ 30 bp. Similarly, specificity depended on cDNA read coverage, read length and sequencing strategy (**Figures 1F-1G**), with >99% specificity achieved in single and paired-end experiments when read length was greater than 50 bp and cDNA coverage ≥5x. In summary, cDNA-detector is highly sensitive and specific even at relatively short read lengths and low contaminant DNA coverage.

### Resource consumption

To test cDNA-detector’s compute resource consumption, we used a human exome alignment (Lim et al., 2015) with 300 M reads. From this exome, we randomly sampled different read numbers and ran cDNA-detector on a Linux high-performance cluster with 72 processors and 251 GB RAM. For small datasets (< 50 M aligned reads), the cDNA detection step was typically completed within 15 minutes and uses ≤ 1 GB memory. cDNA detection on the full exome took about 44 minutes and 2 GB memory (**Figures S3A-S3B**). For large datasets, including whole genome alignments, cDNA-detector can be run in multi-thread mode to decrease processing time, but requires additional memory (**Figures S3C-S3D**). With its low resource footprint, cDNA-detector can easily be integrated into existing NGS processing pipelines as an additional quality control step.

### cDNAs in public databases

We applied cDNA-detector to several large public datasets with different types of high-throughput sequencing experiments to assess potential cDNA cross-contamination. Among 404 ATAC-seq experiments generated from primary tumors in The Cancer Genome Atlas (TCGA; Corces et al., 2018), we identified 16 samples (4%) with suspected exogenous cDNA contamination. Contaminants included the cancer genes *KRAS* (2 samples, including one sample identified by manual review), *STAG2* (8 samples, including one sample identified by manual review), *DDX58* (4 samples; *DDX58* encodes RIG-I), and *SNAP25* (3 samples), some of which can be traced to other projects from the same laboratory (Chen et al., 2017; Mazumdar et al., 2015) (**Figure 2A; Table S1**). Inspection of consensus ATAC-seq peak calls provided with the study revealed that at least some contaminant exons caused spurious peak calls that could affect downstream analyses (**Figure 2A**). Sixteen cDNAs (in 41 samples) were detected in 14,485 ChIP-seq, ATAC-seq and other DNA-seq experiments from the Encyclopedia of DNA Elements (ENCODE) consortium (**Figure 2B; Table S2**). Some of these represent intentionally introduced genes, such as *TERT* for cell line immortalization, or *KLF4* and *MYC* in reprogrammed iPS cells (**Figure S4**). Other genes (*PPARG, PAX7*) could not be traced to intentional transduction in the cell lines in which they were detected and are likely cross-contaminants. Peak calls generated from these experiments and available for public use are affected by these exogenous cDNAs where they contribute signal in unexpected genomic locations (**Figure 2B**). We further performed cDNA detection on a random sampling of 619 human sequencing tracks from 317 projects listed in the NCBI’s Sequence Read Archive (SRA). Fifteen samples with 39 cDNAs were identified in this dataset (**Figure 2C; Table S3**). Detected cDNAs again included intentionally introduced transgenes, but also contaminants that could be traced to other projects in the same laboratory (Huang et al., 2017; Pan et al., 2017; Shi et al., 2019; Yang et al., 2017; **Figure 2C**). A sampling of 71 mouse genome sequencing tracks from 36 projects in SRA yielded three cDNAs. These include intentionally introduced truncated *Sun1* and *NRAS* of human origin (Seehawer et al., 2018), demonstrating that cDNA-detector is mismatch-tolerant and can detect instances of cross-species cDNA contamination (**Figure 2D; Table S4**). No candidate vector-induced cDNAs were detected in 324 exomes from the Cancer Cell Line Encyclopedia (CCLE; Ghandi et al., 2019; data not shown).

**Figure 2:**
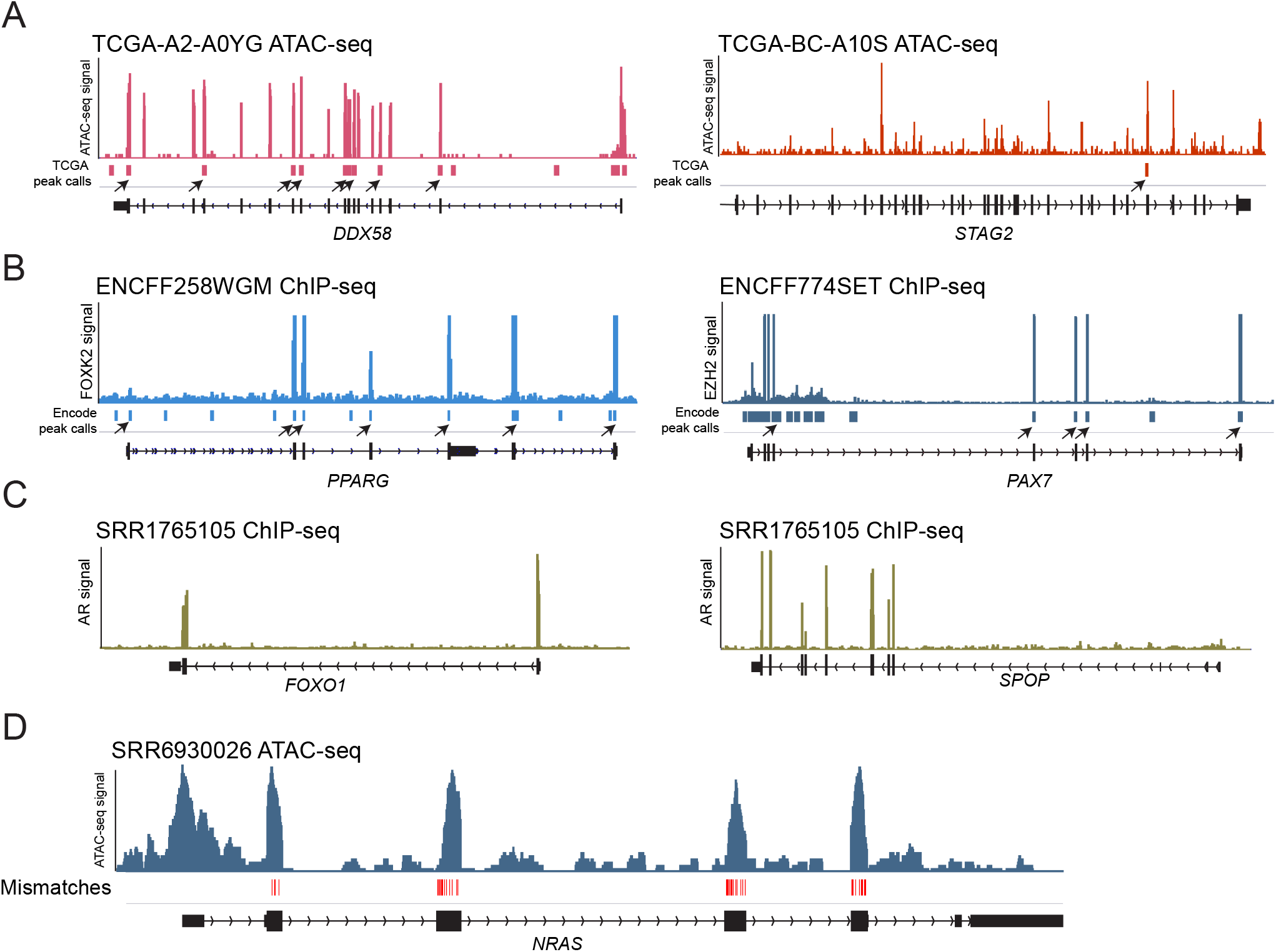
cDNA contamination in published datasets. **A**, Examples of ATAC-seq signal from two primary tumor samples from the TCGA (Corces et al., 2018) showing contamination with *DDX58* (encoding the antiviral innate immune response receptor RIG-I; left) or the cohesin component *STAG2* (right). True signal would be expected at the promoter and potential intragenic regulatory elements, but not over all exons. Arrowheads indicate spurious signal peak calls caused by contaminant reads over exons (black boxes in gene track; peak calls obtained from Corces et al., 2018). **B**, cDNA contamination with *PPARG* in a FOXK2 ChIP-seq experiment in HEK293T cells and *PAX7* in an EZH2 ChIP-seq experiment in HUVEC cells from the ENCODE project. Arrowheads indicate official ENCODE peak calls due to contaminant signal over exons. **C**, Examples of cDNA contamination with prostate cancer genes *FOXO1* and *SPOP* in an androgen receptor (AR) ChIP-seq experiment performed in the prostate cancer cell line C2-4 (Zhao et al., 2016a, 2016b). **D**, Example of transduced human *NRAS* cDNA in a mouse ATAC-seq experiment in cell line ICC2.7.

### Comparison with other methods

Vecuum (Kim et al., 2016) is a similar tool for cDNA detection, also based on evaluating soft-clipped reads at exon boundaries and comparing hits with a static reference vector database. To compare cDNA-detector to Vecuum, we applied cDNA-detector to the human whole-exome sequencing dataset from this publication (Kim et al., 2016; Lim et al., 2015). cDNA-detector detected contamination with *MTOR* cDNA in the same three of the eight exomes (**Figure S5)**. In addition, cDNA-detector discovered *TMEM138* cDNA originating from a vector, used in the same research group (Lee et al., 2012). cDNA-detector further identified additional cDNAs of likely retrogene origin (**Figure S5**). In our NCBI SRA dataset, Vecuum (with lenient parameters) only discovered 16 out of 39 hits found by cDNA-detector (Methods; **Tables S3 and S5**), two out of 41 samples in ENCODE (**Tables S2 and S5**), and two of the hits in the TCGA ATAC-seq dataset. This appears to be caused by short overall read lengths and specifically clipped read length in the ENCODE dataset and a very large number of clipped reads in TCGA ATAC-seq, where adapters are still present in the alignment).

## Discussion

Our results highlight that intentional and unintentional cDNA contamination is pervasive in published sequencing experiments, including some widely used, highly cited datasets, yet remains largely unnoticed. cDNA contaminants are shown to directly affect signal peak calls, and may contribute to other spurious findings. In some cases, specific genes detected as contaminant cDNAs could be traced to other published studies from the submitting laboratory, highlighting the risks of intra-lab contamination. This cross-contamination raises potential issues with keeping specific research constructs confidential and private when necessary. In addition to laboratory protocol improvements, we thus recommend careful review and computational decontamination of high-throughput genomic sequencing data for cDNA contaminants as an essential part of any sequence processing pipeline.

## Methods

### Method outline

The general framework of cDNA-detector consists of three steps: 1. Preparation of a gene model file, for example all exons for a given species and genome assembly, 2. detection of potential cDNA in a BAM-format alignment file, 3. optional clean-up of identified cDNA sequencing reads from the alignment (**Figure 1B; Figures S1 and S2A**).

### Generation of gene model file

An annotation file containing genomic coordinates for exon positions for a given species and assembly is required to run cDNA-detector. A processing script to generate such gene models from GTF format annotation files is part of the cDNA-detector distribution. Pre-generated models for the human (hg19 and hg38) and mouse exome (mm10 and mm39) are provided with the tool. Users may consider adding entries for specific transcript variants (such as truncated genes), custom amplicons, or non-coding RNAs that are suspected or known contaminants.

### Detection of candidate cDNA in DNA sequencing experiments

#### cDNA source reads at exon boundaries

Soft-clipped reads as identified by a read’s CIGAR string (S) in the BAM file are considered potential contaminant reads. In addition, cDNA-detector looks for reads with mismatches just past the exon boundary inside the intron (by MD tag). Reads whose overhang into the intron sequence is an exact substring of the consensus sequence of other soft-clipped reads in this location are also counted as possible cDNA reads. Consensus sequences are generated by aligning the intron overhang of all clipped reads, including a base with ≥ 80% frequency in the consensus.

#### Statistical identification of candidate cDNAs

To evaluate whether the number of soft-clipped boundary-spanning reads for a given exon exceeds the number expected by chance, we build a background expectation model by using all soft-clipped reads overlapping any exon region boundary as defined in the gene model. We then test for significant enrichment of soft-clipped reads, as calculated above, for each single exon boundary genomic position. We define the total number of reads overlapping a specific exon boundary, including genomic reads from the target experiment properly mapped to the adjacent intron, as *n*_*t*_. The total number of soft-clipped and overhang reads is defined as *n*_*c*_. The total number of soft-clipped reads in all exon regions is *N*_*c*_, and the total number of all reads in all exon regions is defined as *N*_*t*_. We use a beta-binomial model to calculate the probability of observing at least *n*_*c*_ reads at a given exon boundary by chance (equation 1):

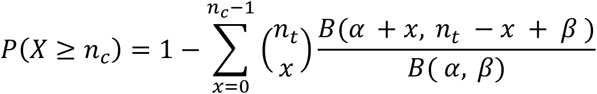

For *X* in {*n*_*c*_, *n*_*c*_ + 1,…, *n*_*t*_}, where α = *N*_*c*_, β = *N*_*t*_ - *N*_*c*_ + 1 and *B*(α, β) is the Beta function. An exon with contaminant cDNA will have soft-clipped reads at both of its boundaries. We therefore combine the *P*-values for both exon boundaries for all exons using the harmonic mean *P*-value (Wilson, 2019) (to account for dependence between the two *P*-values). Finally, all combined exon-wise *P*-values are corrected with the Benjamini-Hochberg procedure (Benjamini and Hochberg, 1995). Exons with corrected *P*-value (*Q*-value) ≤ 0.05 are considered candidate cDNA-exons. To further increase confidence that a flagged exon originates from cDNA, the overhang consensus sequence of soft-clipped reads at both exon boundaries will be compared to the sequence of the neighboring exon(s). If at least one substring match is found, exons are kept for further analysis. Finally, transcripts with ≥ 30% of exons passing the above criteria will be reported.

#### cDNA source inference

cDNA-detector attempts to identify the source of a candidate cDNA to distinguish endogenous (retrocopy) and exogenous (vector) origins. Source inference is performed by querying the overhang sequence of the 5’ and 3’ ends of a candidate cDNA against the NCBI UniVec (https://www.ncbi.nlm.nih.gov/tools/vecscreen/univec/) and Repbase database of human and mouse known repeat sequences (Bao et al., 2015). The highest-scoring match with E-value ≤ 10 is taken as probable origin. Accordingly, a “vector” is reported if the top match stems from UniVec and “retrocopy” is inferred if this hit comes from RepBase. In addition, we require a distance of >5 bp for a repeat match from the clipped position for a “retrocopy” assignment, as these genes often carry adjacent UTR sequence (Casola and Betrán, 2017). In case where no suitable match is found, we use the distance from the exon boundary to the clip as estimate: if the distance is ≤ 5 bp, a cloned vector sequence is more likely (“vector-likely”); if the distance is greater than 10 bp, we infer a possible retrogene (“retrocopy-likely”); otherwise the source is labeled “unknown”. Because source inference strongly depends on the length of the overhang sequence available for BLAST query, it is thus most reliable for experiments with long read lengths.

### Removal of contaminant reads from alignment

After the candidate cDNA contamination detection step, a new alignment file can be generated with candidate contaminant reads removed. This step is straightforward for soft-clipped reads at flagged exon boundaries, as these are very likely contaminants. However, exogenous DNA will also be present as fully aligned reads inside exon regions without overlapping boundaries. In some cases, such as a first exon, these reads can overlap with actual ChIP-seq or ATAC-seq signal derived from the proximal promoter. To address this issue, cDNA-detector assigns reads in candidate cDNA exon regions into three classes: (i) clipped reads and their mates clearly obtained from cDNA contamination (*R*_*c*_); (ii) reads derived from the genome aligned across an exon boundary without soft-clips or mismatches, neighboring exon homology, or a proper mate mapped outside the exon (*R*_*g*_); and (iii) reads which cannot be unambiguously assigned to either class (*R*_*a*_) due to full genomic alignment inside an exon region.

For each candidate contaminant exon, cDNA-detector will first calculate the ratio of *R*_*c*_ to all unambiguous reads (*R*_*c*_ + *R*_*g*_), and then remove the same fraction of ambiguous reads fully mapped inside the exon regions (*R*_*a*_):

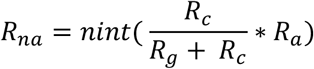

Where *R*_*na*_ is the number of ambiguous reads to remove, *ninit* is the nearest integer function. *R*_*na*_ reads are then randomly selected and removed from this exon. After removal of contaminant reads in all flagged exons, a separate “clean” BAM file is written to a specified output location.

### Performance

#### cDNA simulation experiments for sequencing strategy and read length

To evaluate detection performance, we generated simulated contaminating cDNAs by randomly selecting 100 cDNAs sequences from the CCDS database (Pujar et al., 2018) for each of 10 simulation experiments. Selected cDNAs were “cloned” *in silico* into the pLX307 *vec*tor sequence (http://www.addgene.org/117734/), by replacing the luciferase gene with the corresponding tested cDNA. We then randomly simulated artificial paired-end or single-end reads with read lengths 30 bp, 35 bp, 50 bp, 100 bp, 120 bp, 150 bp, and fragment size 350 bp for paired-end experiments, from the vector with insert, to an average target coverage of 100x. Simulated reads were aligned to the hg38 human genome assembly with bwa mem (Li and Durbin, 2009). Aligned reads were then added to a T47D ATAC-seq experiment with *ERBB2* reads from an intentionally introduced construct removed with cDNA-detector. For experiments with <100x coverage, we randomly downsampled to the desired target coverage from this alignment. cDNA-detector was run with default settings to discover simulated cDNA contamination. Sensitivity is defined as the fraction of detected true positive cDNAs of all 100 inserted cDNAs. Specificity is defined as the fraction of detected true positive cDNAs in all detected cDNAs. Due to possible identical alignments for reads derived from homologous genes, a homolog was also counted as true positive if identified as cDNA candidate by cDNA-detector.

#### Resource consumption

In order to test the time and memory usage and multi-threaded performance of cDNA-detector, we randomly extracted reads to the desired read number from exome SRR1819826 (Lim et al., 2015). Time and memory consumption were assessed for cDNA-detector with default settings.

### Comparison with Vecuum

We directly compared cDNA-detector to Vecuum using the exome data used in the respective publication (SRP055482; Lim et al., 2015). FASTQ files were aligned to the hg19 genome assembly with bwa mem with the default settings (Li and Durbin, 2009). cDNA-detector was run with --num_initial_potential_cdna 5000 (allowing a larger number of initial cDNA candidates for evaluation) to detect cDNA contamination. Candidate cDNAs were manually reviewed for presence of soft-clipped reads and other evidence in the Integrative Genome Viewer (IGV; Robinson et al., 2011).

We also applied Vecuum to the samples from the TCGA, NCBI SRA (human samples only) and ENCODE in which cDNA-detector identified candidate cDNAs. Vecuum run with default parameters (minimal match length *l*=20 and mapping quality threshold *Q=*30) detected few cDNAs in the comparison data. Individual hits could be recovered with Vecuum with more relaxed parameters (-l15 -Q1 for TCGA ATAC-seq; -l 10 -Q1 for NCBI SRA and ENCODE data), although at the cost of lost specificity.

#### Application to public datasets

We applied cDNA-detector snapshot 26 to three large public data sets using the Terra.bio platform and on local compute: 404 ATAC-seq experiments from primary tumors from TCGA (Corces et al., 2018), 14,485 ChIP-seq, ChIA-PET, ATAC-seq and DNase-seq tracks from the ENCODE (Davis et al., 2018) with read length >30 bp, 324 exomes from the Cancer Cell Line Encyclopedia (Ghandi et al., 2019), and 317 human and 36 mouse studies from the NCBI’s SRA repository (see references in **Tables S3 and S4**). SRA studies were selected based on the availability of BAM-format files and restricted to ChIP-seq, ATAC-seq and DNase-seq experiments with sequence read length >30 bp. We then randomly selected one or two (if the study contained more than one experiment) samples from those projects. SRA files were downloaded, then converted to BAM format with sam-dump (https://ncbi.github.io/sra-tools/sam-dump.html) and samtools (Li et al., 2009) with default parameters. Analysis with cDNA-detector was performed as described above. Potential cDNAs were manually reviewed in IGV.

## Acknowledgements

The authors thank Drs. Gad Getz, Michael S. Lawrence and Christopher Ott for their valuable comments on this manuscript. E.R. and M.Q. are supported by funds from the Broad Institute and the MGH Cancer Center. E.R is also supported by funding from the Breast Cancer Alliance.

## Author contributions

M.Q. and E.R. conceived the study. M.Q. wrote the program. U.N., L.S.L. and N.W. provided data and interpretation of results. M.Q. and E.R. analyzed data and wrote the manuscript. All authors have approved the manuscript.

## Conflict of interest statement

The authors declare no conflicting interests.

## Code availability

cDNA-detector is available from GitHub at https://github.com/rheinbaylab/cDNA-detector and as a workflow in Terra.bio.

## Supplementary information

**Table S1:** Identified cDNAs in TCGA ATAC-seq samples

**Table S2:** Identified cDNAs in samples from ENCODE

**Table S3:** Identified cDNAs in select human samples from the NCBI SRA

**Table S4**: Identified cDNAs in select mouse samples from the NCBI SRA

**Table S5:** cDNAs detected by Vecuum in select samples from TCGA, ENCODE and NCBI SRA datasets

## Figure legends

**Figure S1:**
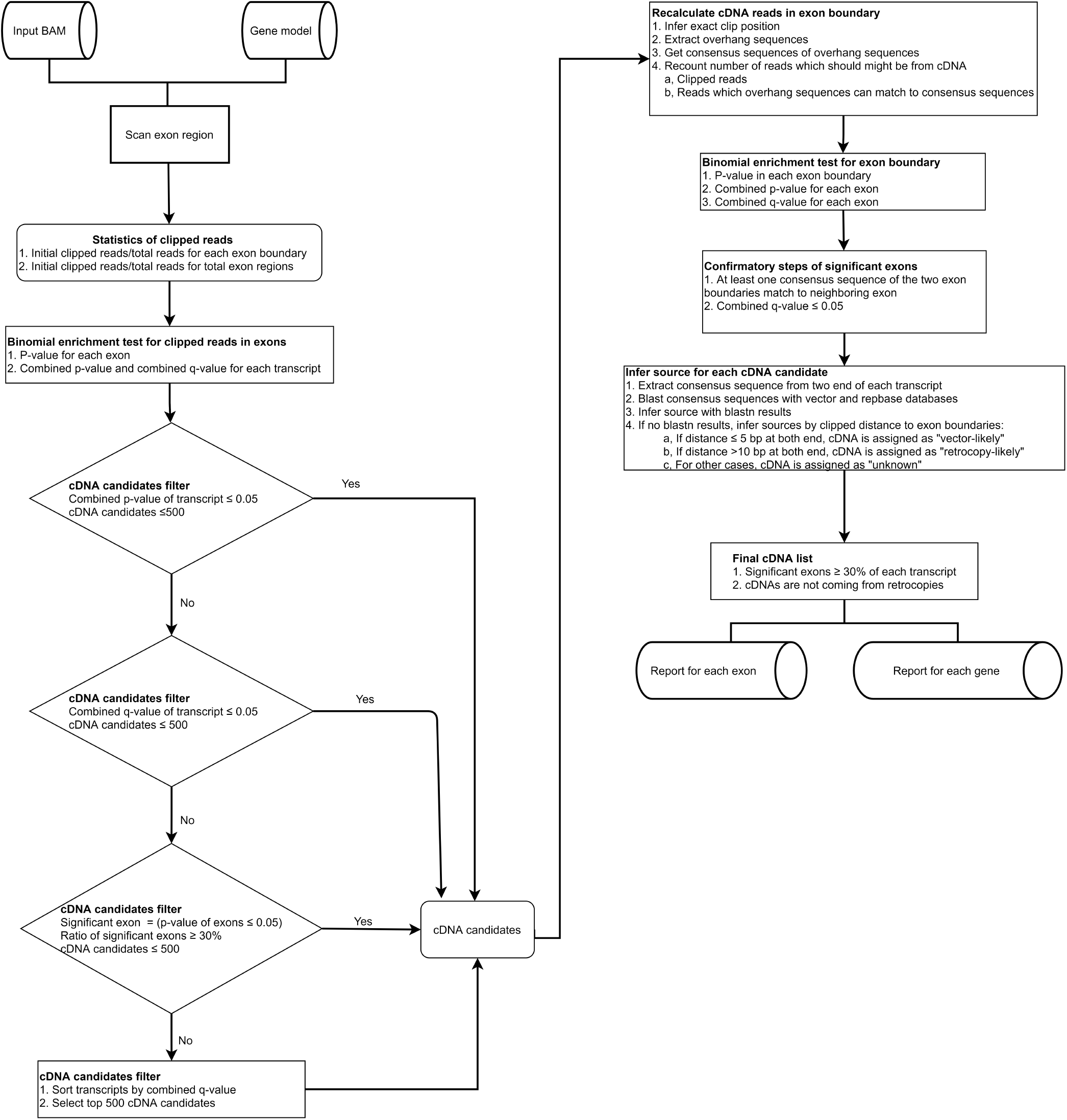
Flowchart detailing the steps of the cDNA-detector detection step.

**Figure S2:**
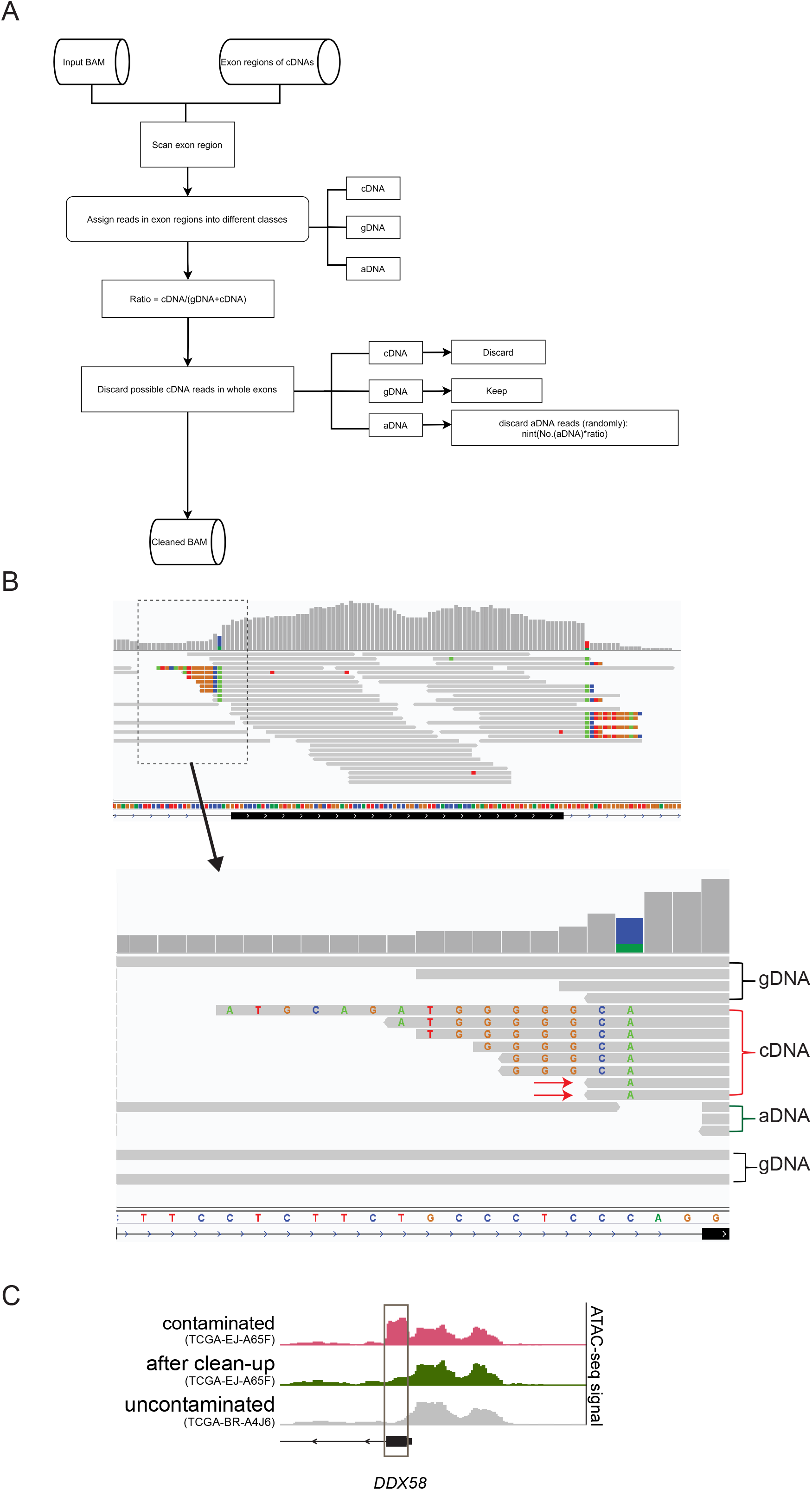
Decontamination of cDNA reads from NGS libraries. **A**, Flowchart depicting the steps of the decontamination step. **B**, Three classes of reads at an exon boundary are considered in the decontamination step: cDNA, genomic DNA (gDNA), and ambiguous reads originating from either cDNA or genomic signal (aDNA). Red arrows show examples of cDNA reads with mismatches instead of clips. **C**, Example of an ATAC-seq experiment contaminated with *DDX58* cDNA and cDNA-detector “decontamination”. Contaminant signal is clearly visible over the first exon, overlapping real ATAC-seq signal from the transcription start site (top, pink track). After the decontamination step, the enrichment over the exon is removed while preserving the ATAC-seq peak (green), comparable to an uncontaminated sample (grey). Gene transcribed from the reverse strand.

**Figure S3:**
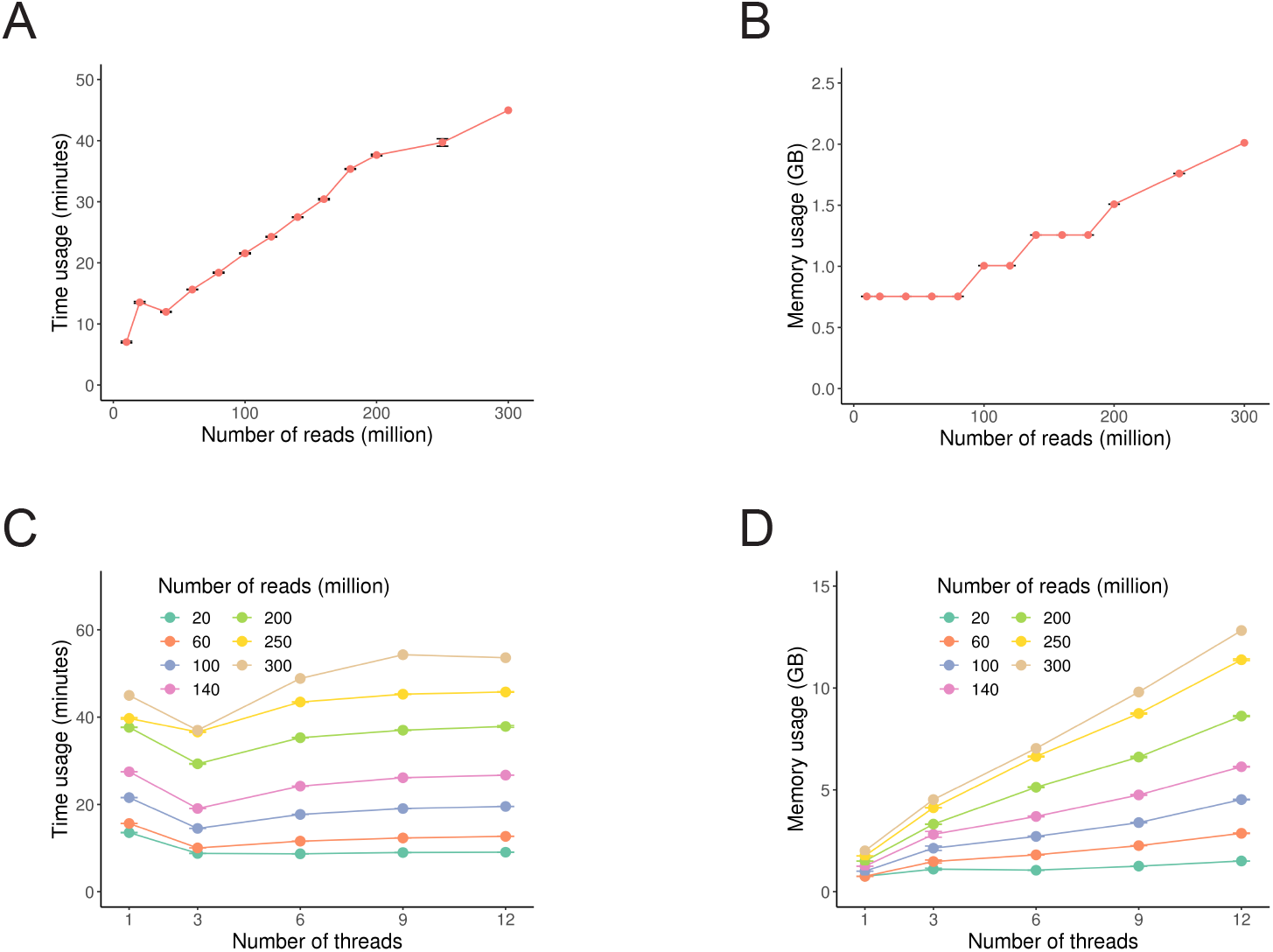
Resource consumption of cDNA-detector. **A**, Runtime of the cDNA-detector detection step depending on the number of reads in the alignment. **B**, Memory usage in GB of the detection step for different number of reads. **C-D**: Time (**C**) and memory (**D**) usage for different read numbers when using multiple threads. The error bars indicate standard errors. Sample size n = 10 for each simulation (except the full data set of 300 M reads).

**Figure S4:**
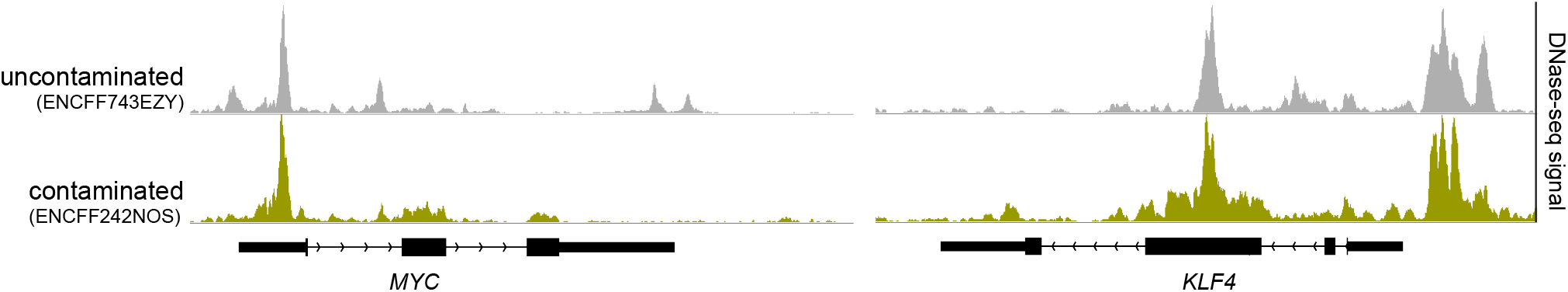
cDNA identified in a reprogrammed iPS cell line. *MYC* and *KLF4* cDNA in L1-S8R cells (yellow) reprogrammed with *OCT4, SOX2, KLF4, cMYC*, and uncontaminated control cell line PC-3 (grey).

**Figure S5:**
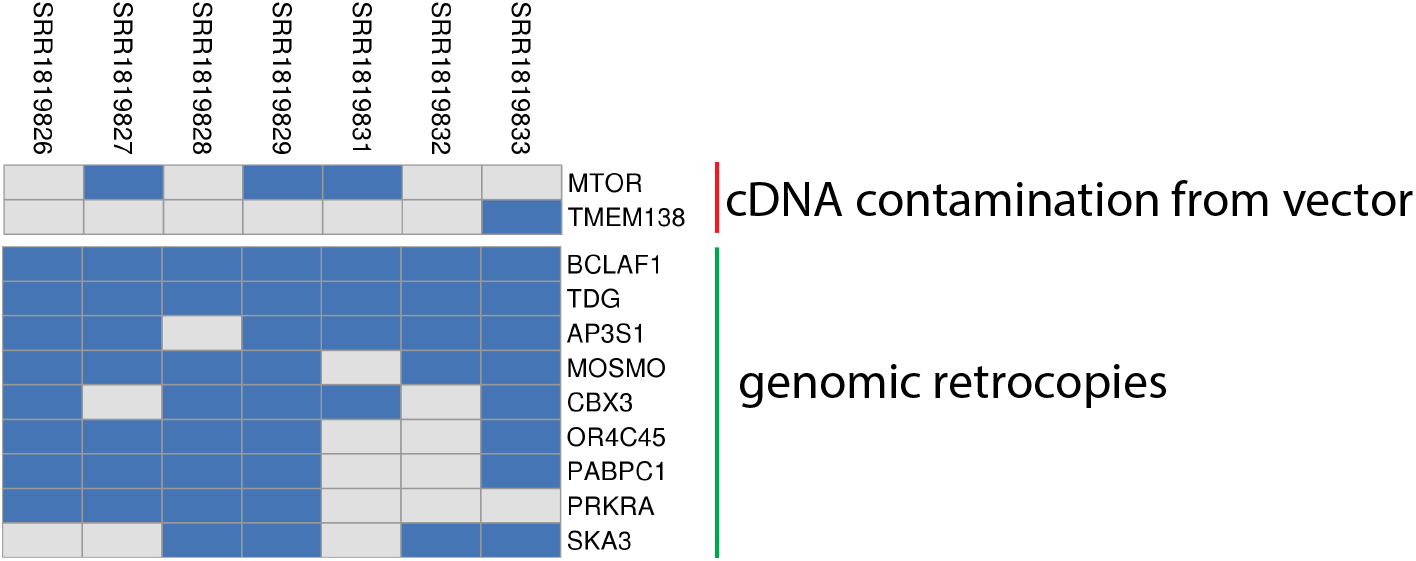
cDNA identified by cDNA-detector in a human whole-exome sequencing dataset. Identification of known vector contamination (*MTOR*), previously unknown vector contamination (*TMEM138*) and genomic retrocopies of processed genes by cDNA-detector in the dataset presented in the Vecuum publication (Kim et al., 2016).

## Notes

### Competing Interest Statement

The authors have declared no competing interest.

## References

Bao, W., Kojima, K.K., and Kohany, O. (2015). Repbase Update, a database of repetitive elements in eukaryotic genomes. Mob. DNA 6, 11.

Benjamini, Y., and Hochberg, Y. (1995). Controlling the false discovery rate: A practical and powerful approach to multiple testing. J. R. Stat. Soc. 57, 289–300.

Casola, C., and Betrán, E. (2017). The Genomic Impact of Gene Retrocopies: What Have We Learned from Comparative Genomics, Population Genomics, and Transcriptomic Analyses? Genome Biol. Evol. 9, 1351–1373.

Chen, Y.G., Kim, M.V., Chen, X., Batista, P.J., Aoyama, S., Wilusz, J.E., Iwasaki, A., and Chang, H.Y. (2017). Sensing Self and Foreign Circular RNAs by Intron Identity. Mol. Cell 67, 228–238.e5.

Corces, M.R., Granja, J.M., Shams, S., Louie, B.H., Seoane, J.A., Zhou, W., Silva, T.C., Groeneveld, C., Wong, C.K., Cho, S.W., et al. (2018). The chromatin accessibility landscape of primary human cancers. Science 362.

Davis, C.A., Hitz, B.C., Sloan, C.A., Chan, E.T., Davidson, J.M., Gabdank, I., Hilton, J.A., Jain, K., Baymuradov, U.K., Narayanan, A.K., et al. (2018). The Encyclopedia of DNA elements (ENCODE): data portal update. Nucleic Acids Res. 46, D794–D801.

Esnault, C., Maestre, J., and Heidmann, T. (2000). Human LINE retrotransposons generate processed pseudogenes. Nat. Genet. 24, 363–367.

Ghandi, M., Huang, F.W., Jané-Valbuena, J., Kryukov, G.V., Lo, C.C., McDonald, E.R., 3rd, Barretina, J., Gelfand, E.T., Bielski, C.M., Li, H., et al. (2019). Next-generation characterization of the Cancer Cell Line Encyclopedia. Nature 569, 503–508.

Huang, S.N., Williams, J.S., Arana, M.E., Kunkel, T.A., and Pommier, Y. (2017). Topoisomerase I-mediated cleavage at unrepaired ribonucleotides generates DNA double-strand breaks. EMBO J. 36, 361–373.

Kaessmann, H., Vinckenbosch, N., and Long, M. (2009). RNA-based gene duplication: mechanistic and evolutionary insights. Nat. Rev. Genet. 10, 19–31.

Kim, J., Maeng, J.H., Lim, J.S., Son, H., Lee, J., Lee, J.H., and Kim, S. (2016). Vecuum: identification and filtration of false somatic variants caused by recombinant vector contamination. Bioinformatics 32, 3072–3080.

Kim, J., Zhao, B., Huang, A.Y., Miller, M.B., Lodato, M.A., Walsh, C.A., and Lee, E.A. (2020). APP gene copy number changes reflect exogenous contamination. Nature 584, E20–E28.

Lee, J.H., Silhavy, J.L., Lee, J.E., Al-Gazali, L., Thomas, S., Davis, E.E., Bielas, S.L., Hill, K.J., Iannicelli, M., Brancati, F., et al. (2012). Evolutionarily assembled cis-regulatory module at a human ciliopathy locus. Science 335, 966–969.

Lee, M.-H., Siddoway, B., Kaeser, G.E., Segota, I., Rivera, R., Romanow, W.J., Liu, C.S., Park, C., Kennedy, G., Long, T., et al. (2018). Somatic APP gene recombination in Alzheimer’s disease and normal neurons. Nature 563, 639–645.

Li, H., and Durbin, R. (2009). Fast and accurate short read alignment with Burrows-Wheeler transform. Bioinformatics 25, 1754–1760.

Li, H., Handsaker, B., Wysoker, A., Fennell, T., Ruan, J., Homer, N., Marth, G., Abecasis, G., Durbin, R., and 1000 Genome Project Data Processing Subgroup (2009). The Sequence Alignment/Map format and SAMtools. Bioinformatics 25, 2078–2079.

Lim, J.S., Kim, W.-I., Kang, H.-C., Kim, S.H., Park, A.H., Park, E.K., Cho, Y.-W., Kim, S., Kim, H.M., Kim, J.A., et al. (2015). Brain somatic mutations in MTOR cause focal cortical dysplasia type II leading to intractable epilepsy. Nat. Med. 21, 395–400.

Mazumdar, C., Shen, Y., Xavy, S., Zhao, F., Reinisch, A., Li, R., Corces, M.R., Flynn, R.A., Buenrostro, J.D., Chan, S.M., et al. (2015). Leukemia-Associated Cohesin Mutants Dominantly Enforce Stem Cell Programs and Impair Human Hematopoietic Progenitor Differentiation. Cell Stem Cell 17, 675–688.

Pan, C.-W., Jin, X., Zhao, Y., Pan, Y., Yang, J., Karnes, R.J., Zhang, J., Wang, L., and Huang, H. (2017). AKT-phosphorylated FOXO1 suppresses ERK activation and chemoresistance by disrupting IQGAP1-MAPK interaction. EMBO J. 36, 995–1010.

Pujar, S., O’Leary, N.A., Farrell, C.M., Loveland, J.E., Mudge, J.M., Wallin, C., Girón, C.G., Diekhans, M., Barnes, I., Bennett, R., et al. (2018). Consensus coding sequence (CCDS) database: a standardized set of human and mouse protein-coding regions supported by expert curation. Nucleic Acids Res. 46, D221–D228.

Robinson, J.T., Thorvaldsdóttir, H., Winckler, W., Guttman, M., Lander, E.S., Getz, G., and Mesirov, J.P. (2011). Integrative genomics viewer. Nat. Biotechnol. 29, 24–26.

Seehawer, M., Heinzmann, F., D’Artista, L., Harbig, J., Roux, P.-F., Hoenicke, L., Dang, H., Klotz, S., Robinson, L., Doré, G., et al. (2018). Necroptosis microenvironment directs lineage commitment in liver cancer. Nature 562, 69–75.

Shi, Q., Zhu, Y., Ma, J., Chang, K., Ding, D., Bai, Y., Gao, K., Zhang, P., Mo, R., Feng, K., et al. (2019). Prostate Cancer-associated SPOP mutations enhance cancer cell survival and docetaxel resistance by upregulating Caprin1-dependent stress granule assembly. Molecular Cancer 18.

Wei, W., Gilbert, N., Ooi, S.L., Lawler, J.F., Ostertag, E.M., Kazazian, H.H., Boeke, J.D., and Moran, J.V. (2001). Human L1 retrotransposition: cis preference versus trans complementation. Mol. Cell. Biol. 21, 1429–1439.

Wilson, D.J. (2019). The harmonic mean p-value for combining dependent tests. Proceedings of the National Academy of Sciences. 116, 1195–1200.

Yang, Y., Blee, A.M., Wang, D., An, J., Pan, Y., Yan, Y., Ma, T., He, Y., Dugdale, J., Hou, X., et al. (2017). Loss of FOXO1 Cooperates with TMPRSS2–ERG Overexpression to Promote Prostate Tumorigenesis and Cell Invasion. Cancer Res. 77, 6524–6537.

Zhao, J., Zhao, Y., Wang, L., Zhang, J., Karnes, R.J., Kohli, M., Wang, G., and Huang, H. (2016a). Alterations of androgen receptor-regulated enhancer RNAs (eRNAs) contribute to enzalutamide resistance in castration-resistant prostate cancer. Oncotarget 7, 38551–38565.

Zhao, Y., Wang, L., Ren, S., Wang, L., Blackburn, P.R., McNulty, M.S., Gao, X., Qiao, M., Vessella, R.L., Kohli, M., et al. (2016b). Activation of P-TEFb by Androgen Receptor-Regulated Enhancer RNAs in Castration-Resistant Prostate Cancer. Cell Rep. 15, 599–610.

